## Supplementary figures and images for "The intrinsically disordered N-terminus of the voltage-dependent anion channel"

### S1 Fig

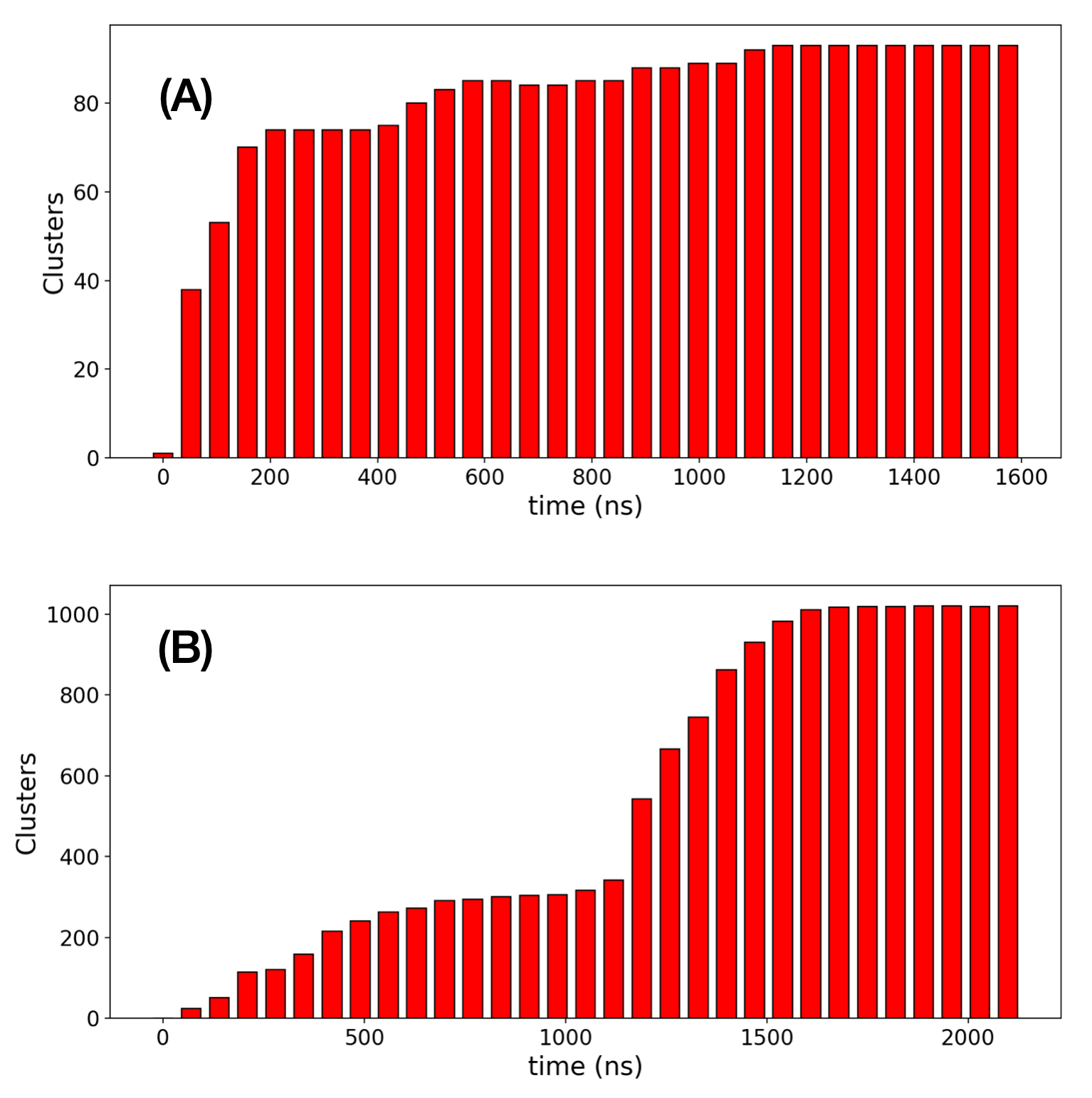

### S1 Table

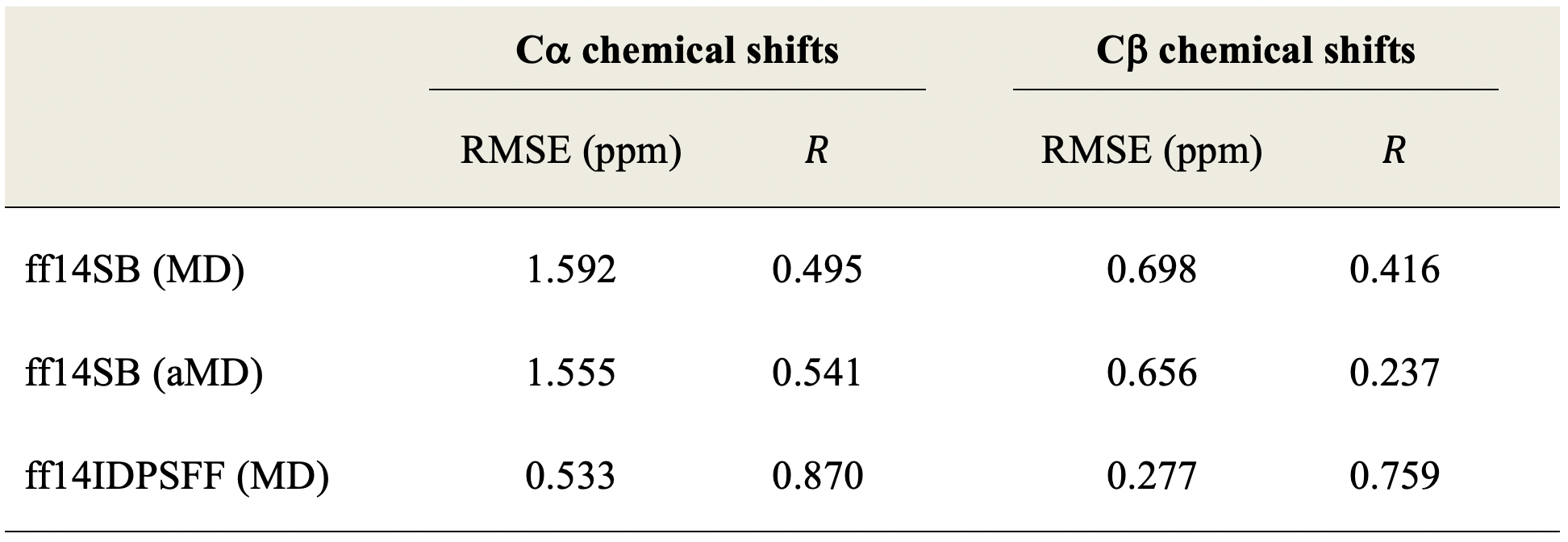

### S2 Fig

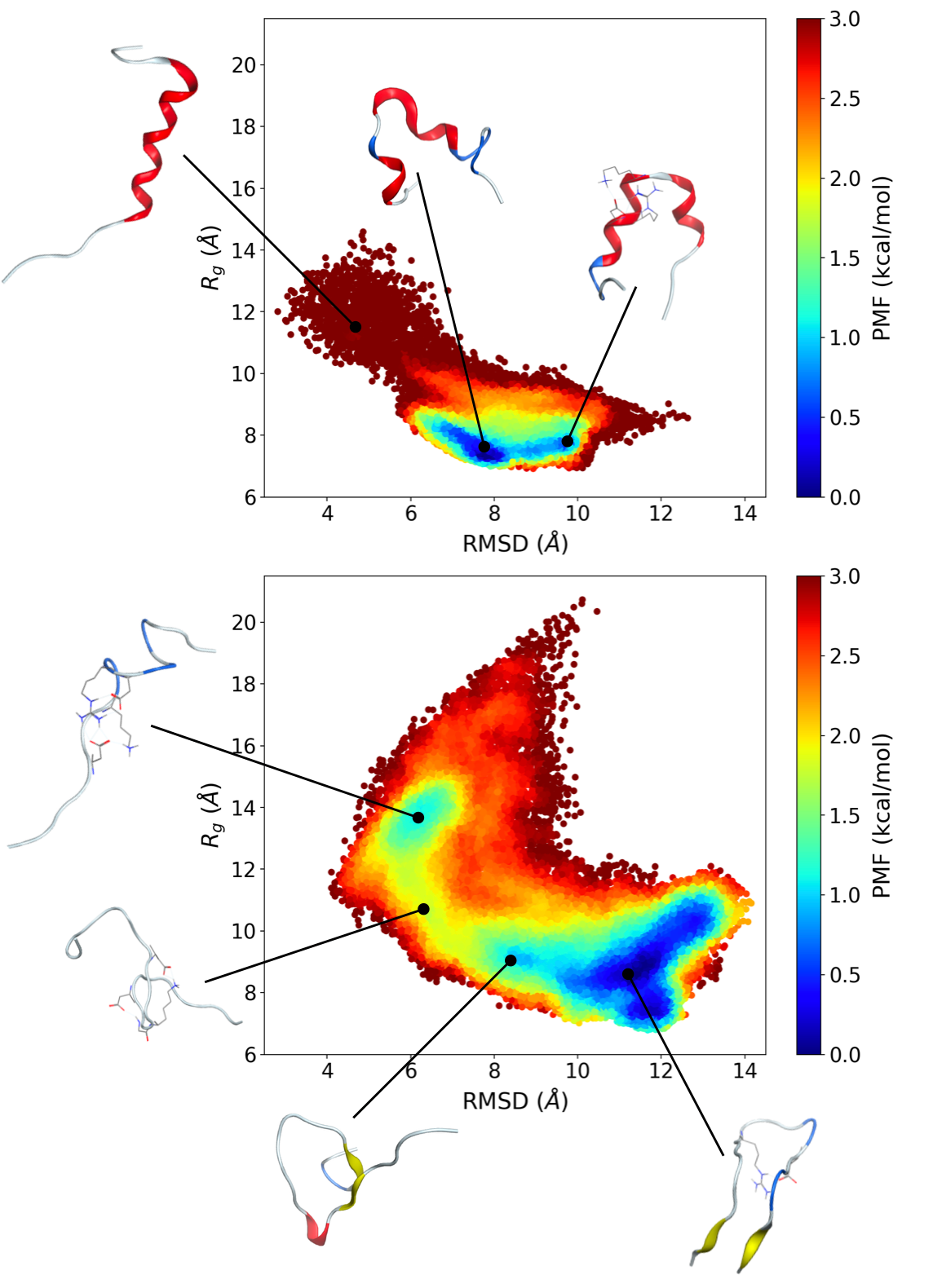

### S2 Table

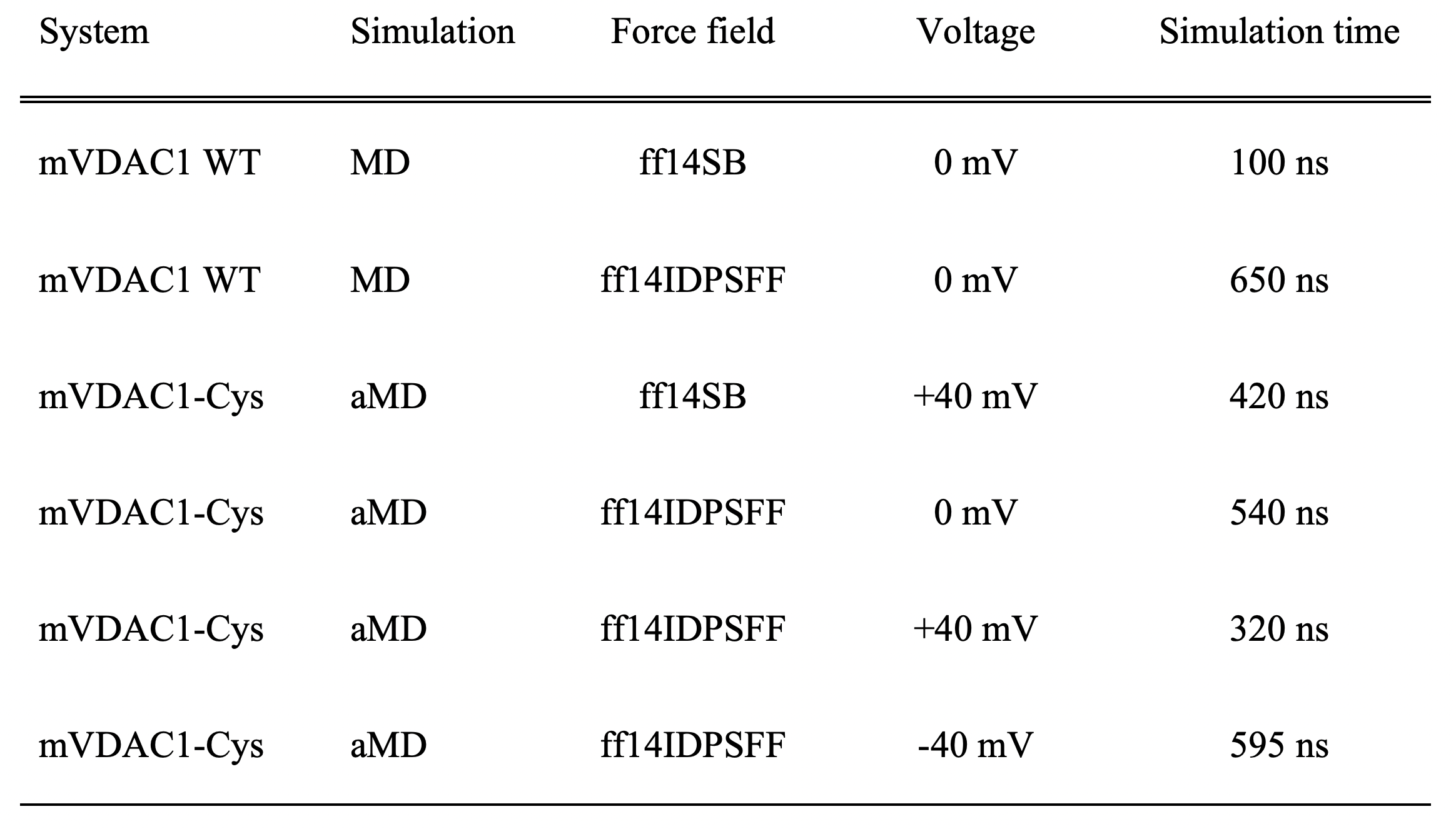

### S3 Fig

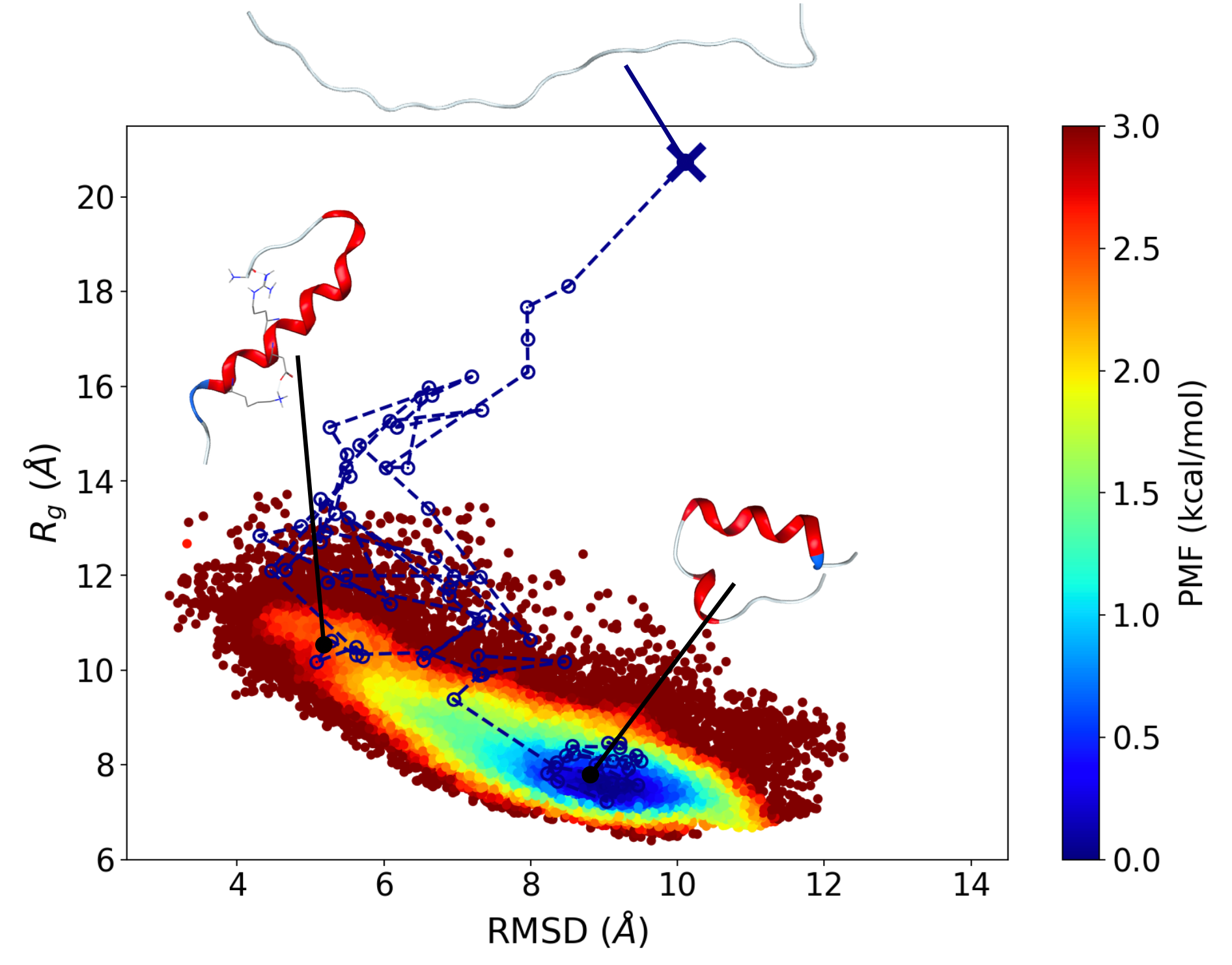

### S3 Table

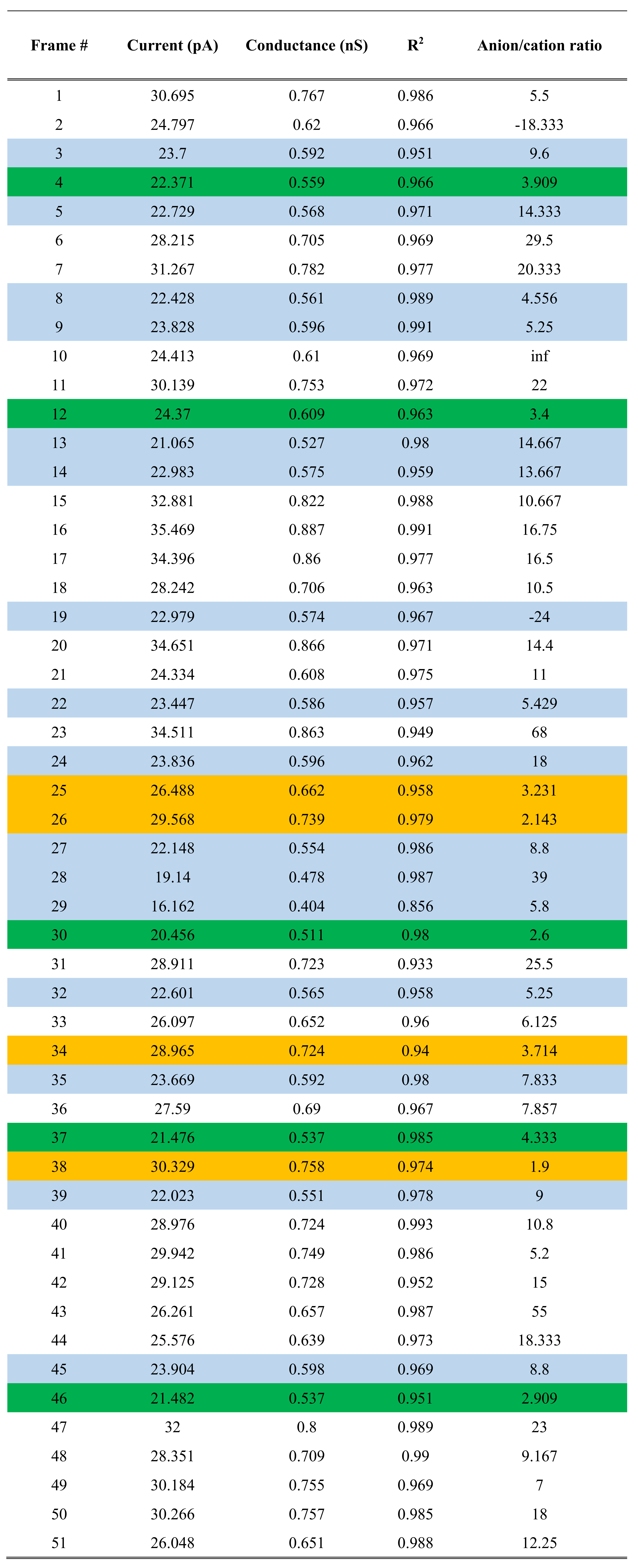

### S4 Fig

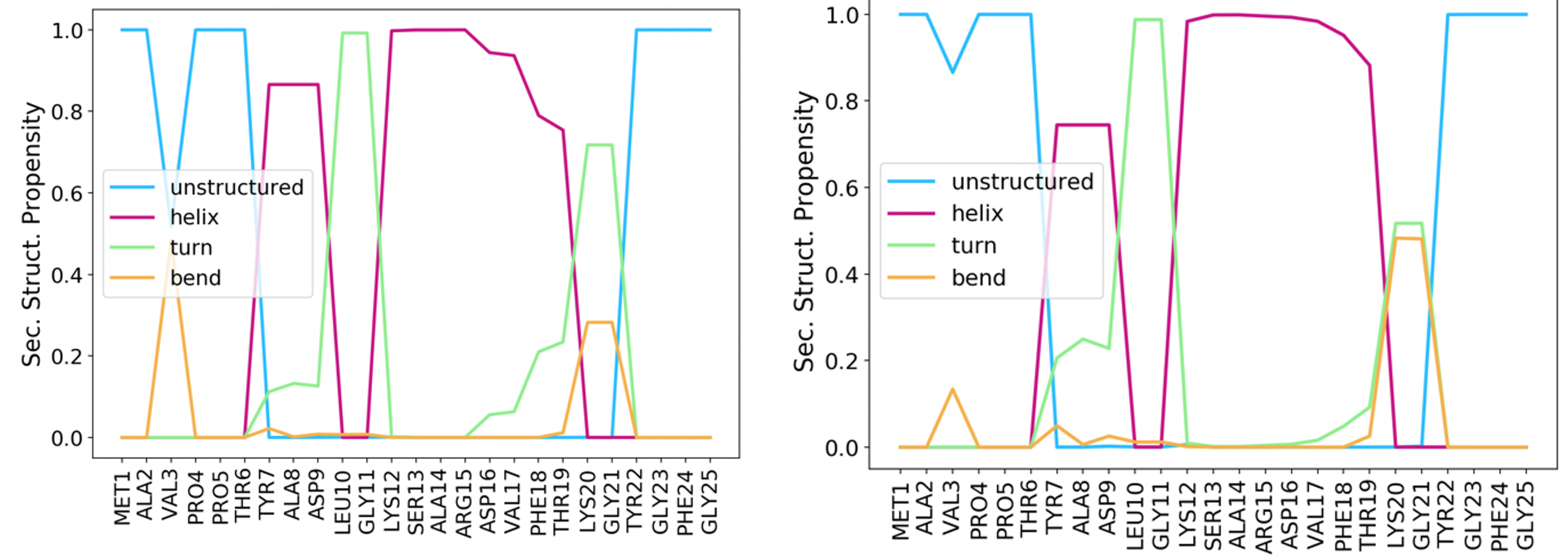

### S5 Fig

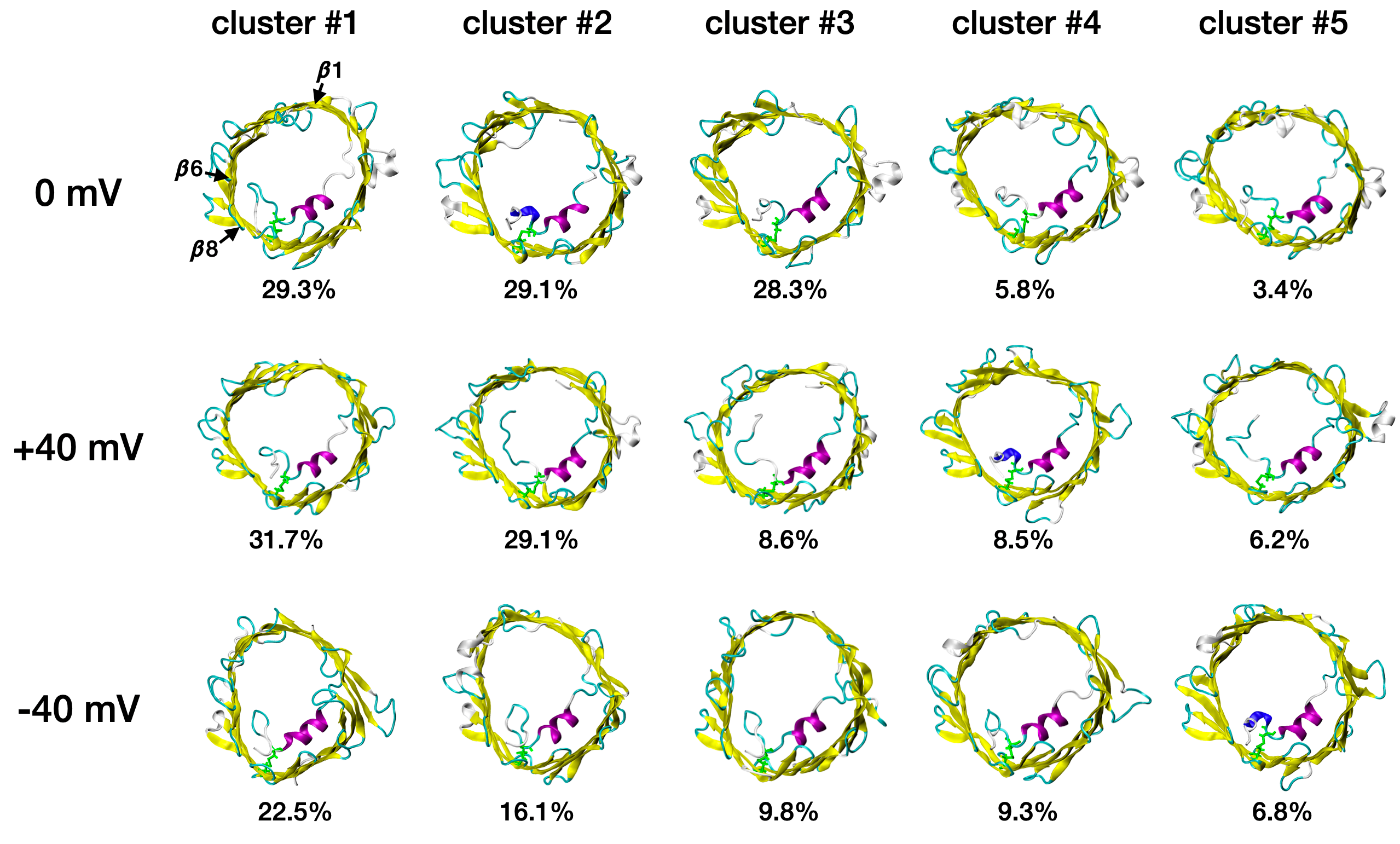

### S6 Fig

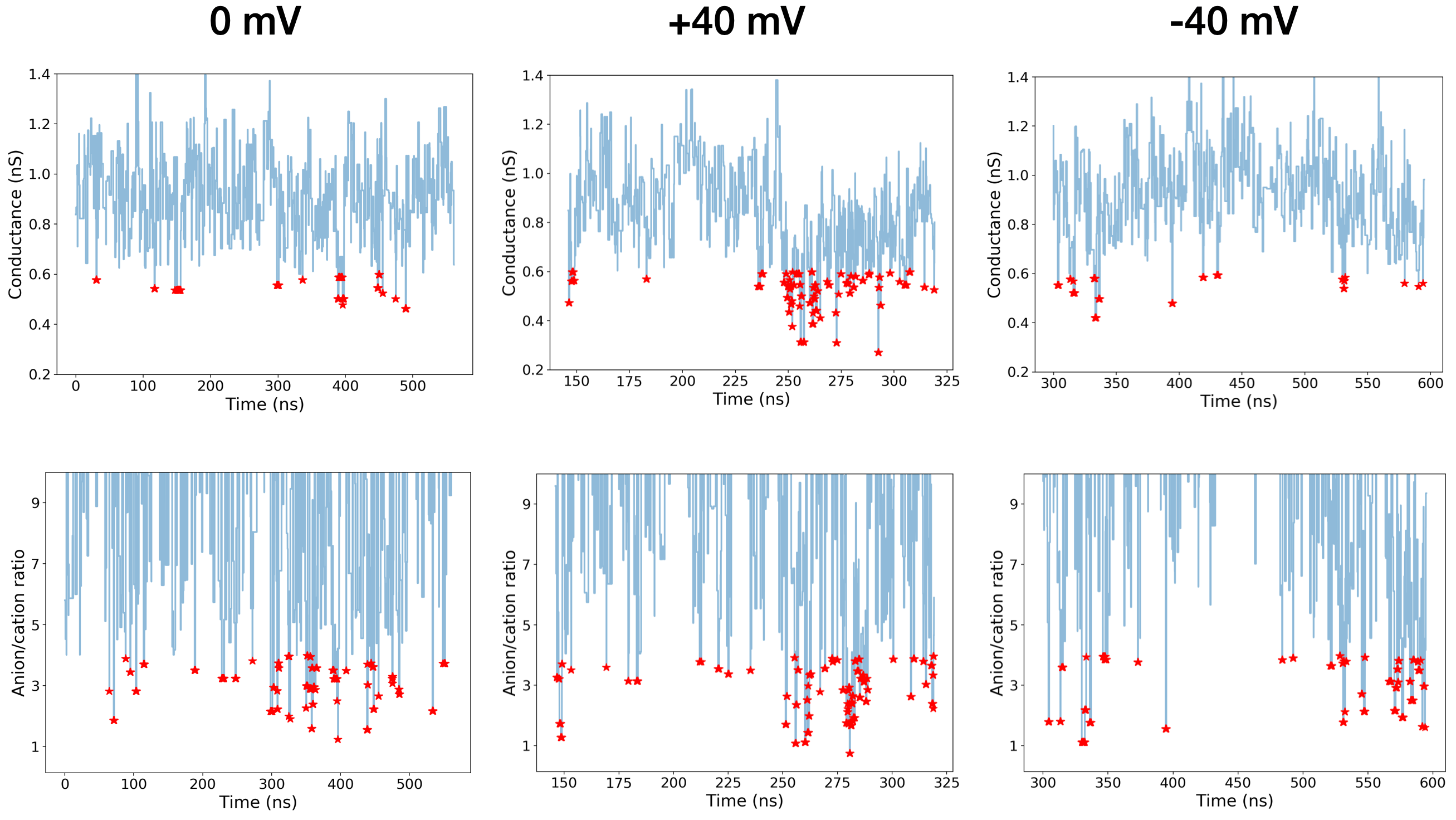

### S7 Fig

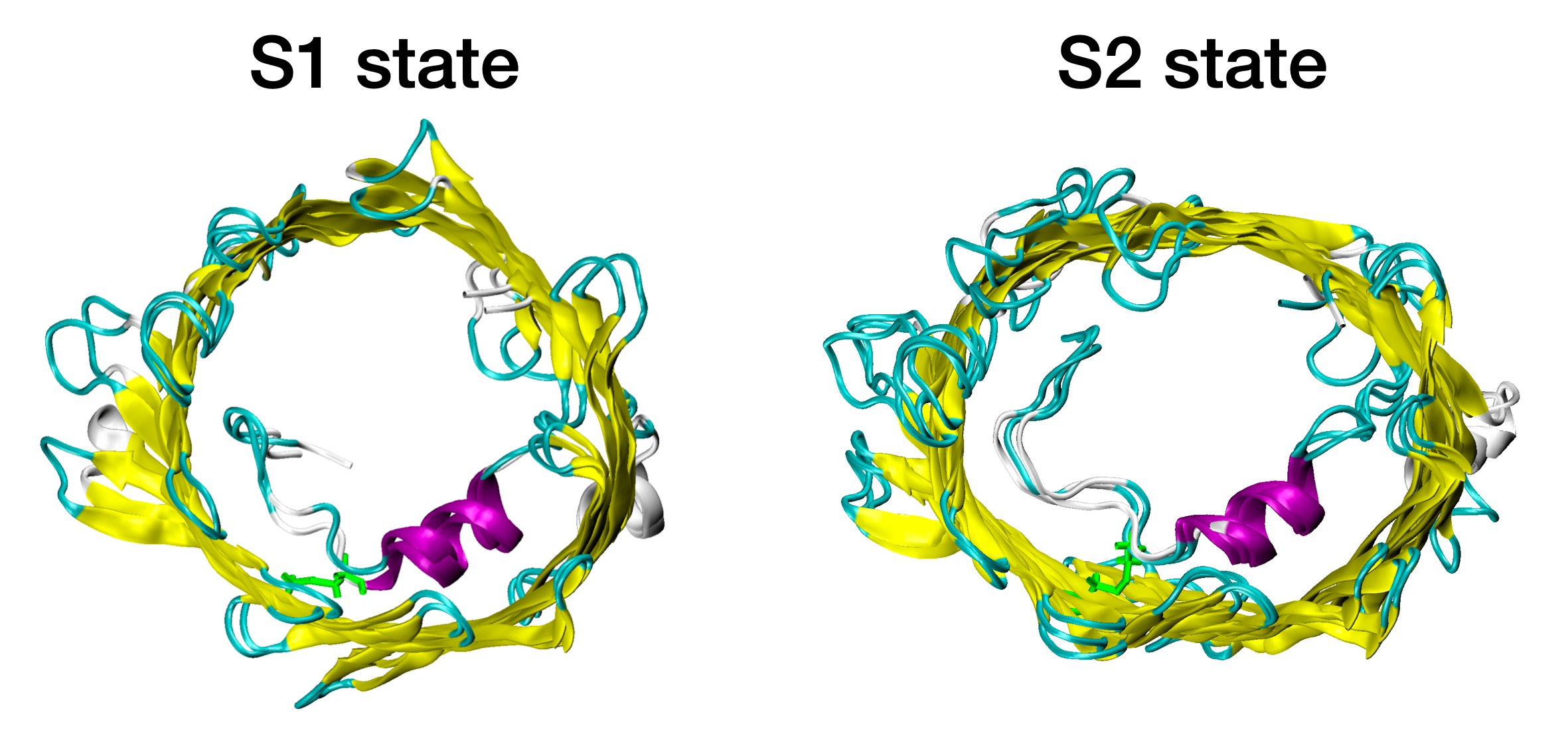

### S8 Fig

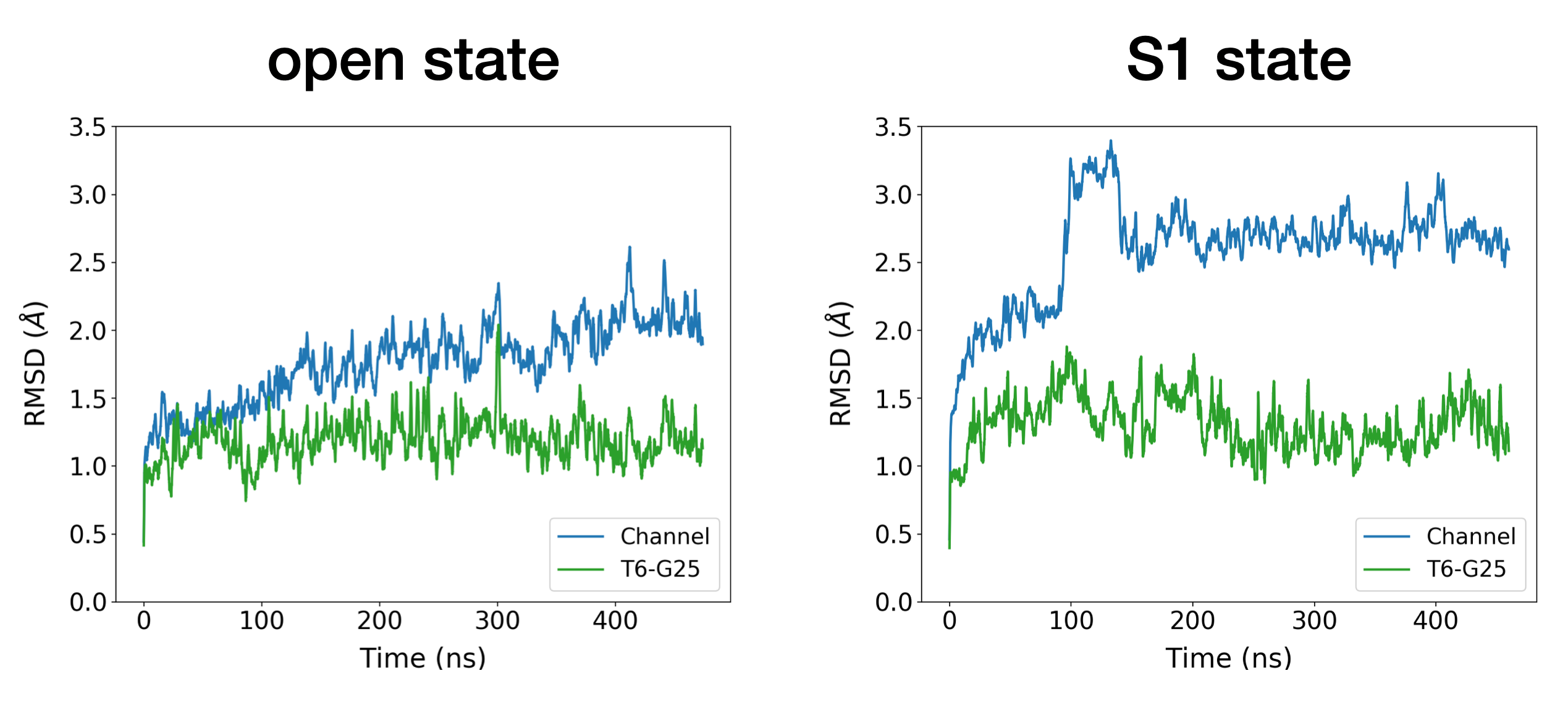

### S9 Fig

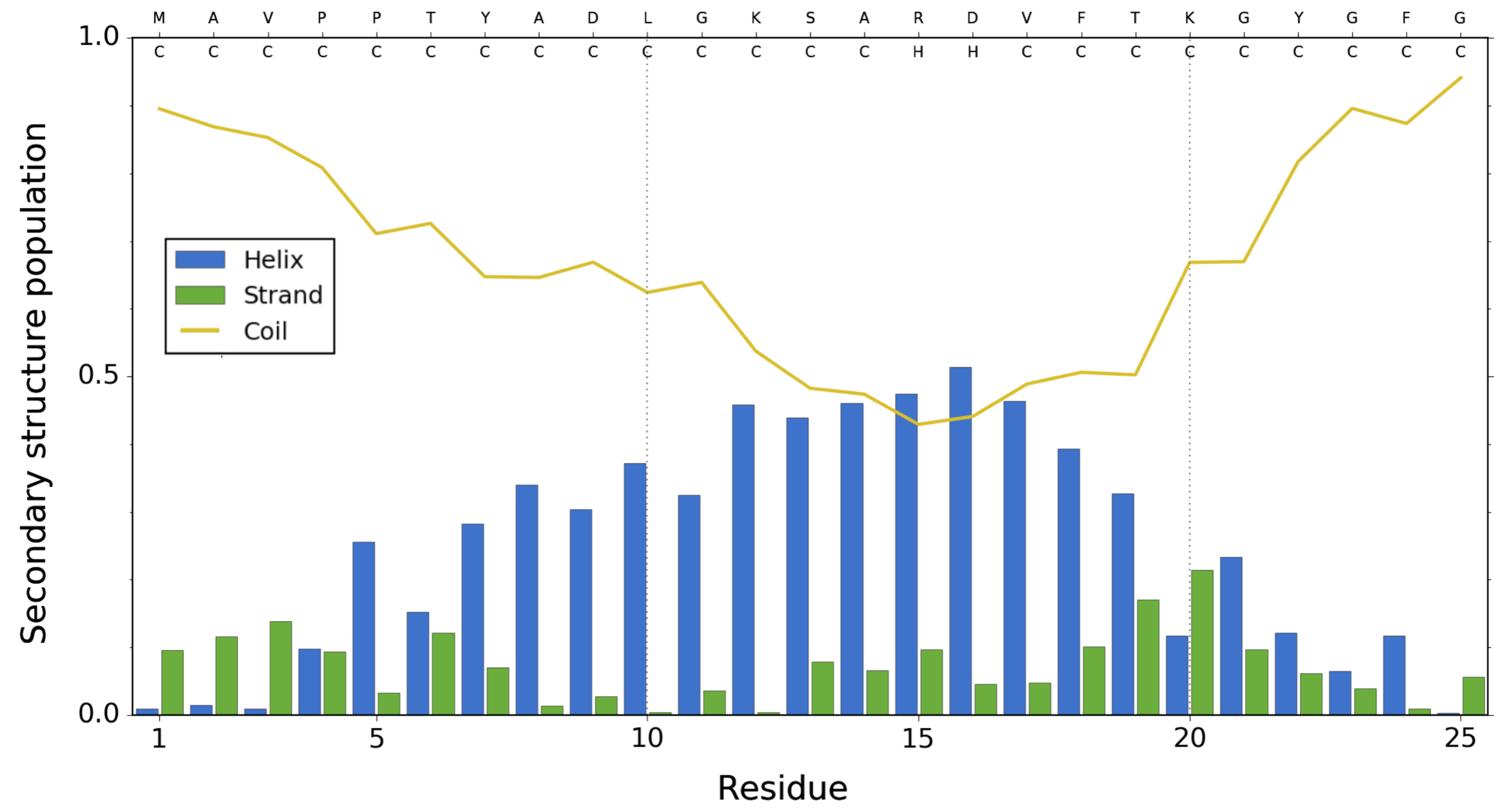

### S10 Fig

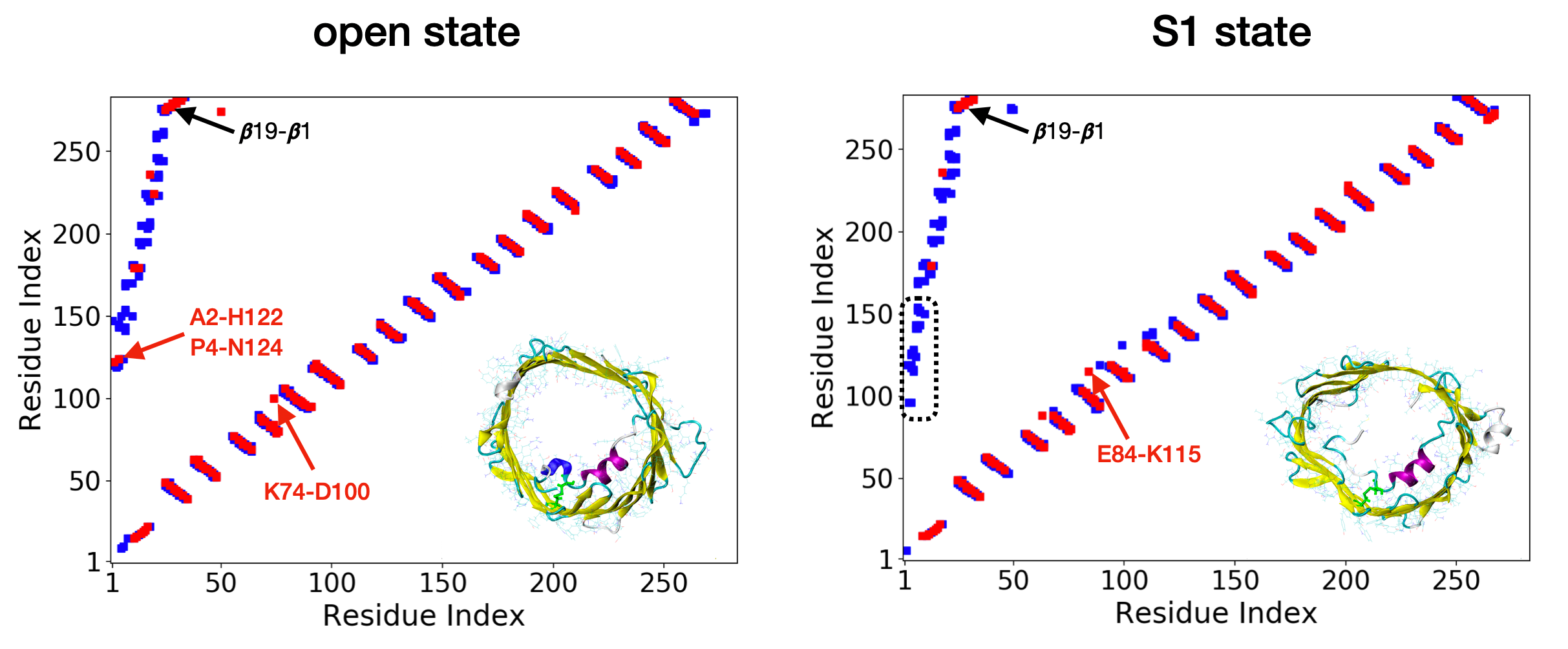
